# Towards VDAC pharmacology: Developing small-peptide channel inhibitors inspired by nature

**DOI:** 10.64898/2026.10.13.758907

**Authors:** Tatiana K. Rostovtseva, Philip A. Gurnev, David P. Hoogerheide, Sergey M. Bezrukov

**Affiliations:** Section on Molecular Transport, Eunice Kennedy Shriver National Institute of Child Health and Human Development, NIH, Bethesda, MD; Center for Neutron Research, National Institute of Standards and Technology, Gaithersburg, MD

**Keywords:** protein-nanopore interactions, peripheral proteins, mitochondrial membranes, energy landscape modeling

## Abstract

The Voltage-Dependent Anion Channel (VDAC) forms a unique pathway across the mitochondrial outer membrane for small ions and water-soluble metabolites, such as ATP and ADP. Despite VDAC’s involvement in both normal regulation of respiration and in a wide variety of mitochondria-associated pathologies, there are, surprisingly, no known FDA-approved pharmacological agents targeting VDAC. Earlier, we have shown that VDAC can be regulated through its interactions with well-known cytosolic proteins – dimeric tubulin and α-synuclein (αSyn) – that are able to reversibly block this channel at nanomolar concentrations. We hypothesized that the architecture of a prospective VDAC inhibitor should fulfill two main requirements: possess a negatively charged disordered polypeptide C- or N-terminus as a pore blocking domain and feature a mitochondrial membrane-binding domain. Using αSyn as a template, we engineered three peptides and demonstrated that they all characteristically block VDAC reconstituted into planar lipid membranes in a controllable manner, thereby mimicking the actions of tubulin and αSyn. The VDAC blockage times induced by the synthetic peptides are comparable to αSyn-induced blockages, though the half-effect solution concentration for synthetic blockers is a hundred times higher than that of αSyn. Our work can be considered a proof-of-principle study that informs future development of mitochondria-targeting pharmacological tools.

## Introduction

About 15-20% of all marketed drugs act as modifiers of ion channels, generating over ten billion US$ annually in worldwide sales ^1-4^. These channel-targeting drugs are indispensable in modern health care. However, to date, this progress has been mostly limited to ion-selective channels of neurophysiology and toxin channels ^5-12^. Surprisingly, there are zero known specifically designed and FDA-approved pharmacological agents for the voltage-dependent anion channel (VDAC) of the mitochondrial outer membrane (MOM), despite this channel’s pivotal role in both normal respiration regulation in health and development, as well as in a wide variety of mitochondria-associated pathologies. VDAC has been shown to be implicated in diabetes, hypertension, cardiovascular diseases, various cancers, and neurodegenerative disorders, such as Parkinson’s, Alzheimer’s, and amyotrophic lateral sclerosis ^13^.

Due to its central role in controlling MOM permeability and, consequently, mitochondrial function ^14, 15^, VDAC is emerging as a promising pharmacological target for treating a wide variety of mitochondria-associated pathologies ^16, 17^. Proteomics and biochemical analyses have identified a wide range of non-polar compounds, including natural and synthetic steroids, as well as small-molecule anticancer and neuroprotective drugs, that bind to VDAC ^18-20^. Therefore, understanding the molecular mechanism of VDAC functioning and identifying both its natural regulators and synthetic inhibitors is vital for improving outcomes of mitochondria-associated diseases and for utilizing VDAC as a promising pharmacological target. However, progress in this direction is still painfully slow due to the absence of a potent VDAC inhibitor that, among other applications, could be a long-awaited tool for a wide research community to selectively block VDAC in cells.

Although recognizing that the road to a successful design and practical implementation of VDAC-targeting drugs is long and challenging, we share below what we think are promising results and ideas that point in the right direction. Our efforts are inspired by our discovery of a naturally occurring mechanism of VDAC regulation by cytosolic proteins. Specifically, we found that VDAC is dynamically controlled by its interactions with cytosolic proteins that possess negatively charged disordered termini. We have identified two such proteins – dimeric tubulin and α-synuclein (αSyn) – as highly potent regulators of VDAC, able to inhibit this channel with nanomolar efficiency ^21, 22^ (M = mol/L).

Our previous work has led to a detailed understanding of the molecular mechanisms by which these cytosolic proteins interact with mitochondrial membranes and VDAC. We were able to assign specific roles to the dimeric tubulin globular body and its C-terminus, with the former serving as a membrane anchor and the latter as a pore blocker. Similarly, both the N- and C-termini of αSyn were shown to be involved in its blockage of VDAC. The C-terminus enters the net positive VDAC pore and partially blocks VDAC, while the N-terminus binds to the membrane, preventing αSyn translocation. As we confirmed in experiments with MspA and α-hemolysin, another two β-barrel channels, the N-terminus of αSyn becomes detached from the membrane surface only at relatively high applied potentials, allowing the whole molecule to translocate through the channel in in vitro experiments ^23^.

We thus proposed a model of VDAC inhibition that explains how these two very different and otherwise unrelated abundant cytosolic proteins induce qualitatively similar blockages of VDAC ^24^. There are several important similarities: both proteins have a disordered, highly negatively charged C-terminus and are “amphitropic” or peripheral, i.e., they are water-soluble and reversibly associate with cellular membranes ^25^. Specifically, they are able to bind to mitochondrial membranes in live cells ^26-29^ and modulate mitochondrial metabolism and respiration by regulating VDAC permeability in a lipid-dependent manner ^30-32^. Generalizing from these two natural examples, we hypothesized that the architecture of a VDAC inhibitor should fulfill two main requirements: possess a negatively charged, disordered C- or N-terminus and a membrane-binding domain ^24^. Importantly, the proposed model of VDAC inhibition is quite general, does not require any specific interaction between the VDAC pore and inhibitor, and suggests that other cytosolic proteins or synthetic peptides may also be involved. The model is illustrated in **Fig. 1**. According to it, the process of VDAC interaction with a cytosolic inhibitor consists of three steps: the inhibitor adsorption to the membrane surface, lateral diffusion to the channel, and voltage-driven capture of the charged terminus of the inhibitor protein by the VDAC pore.

**Fig. 1.**
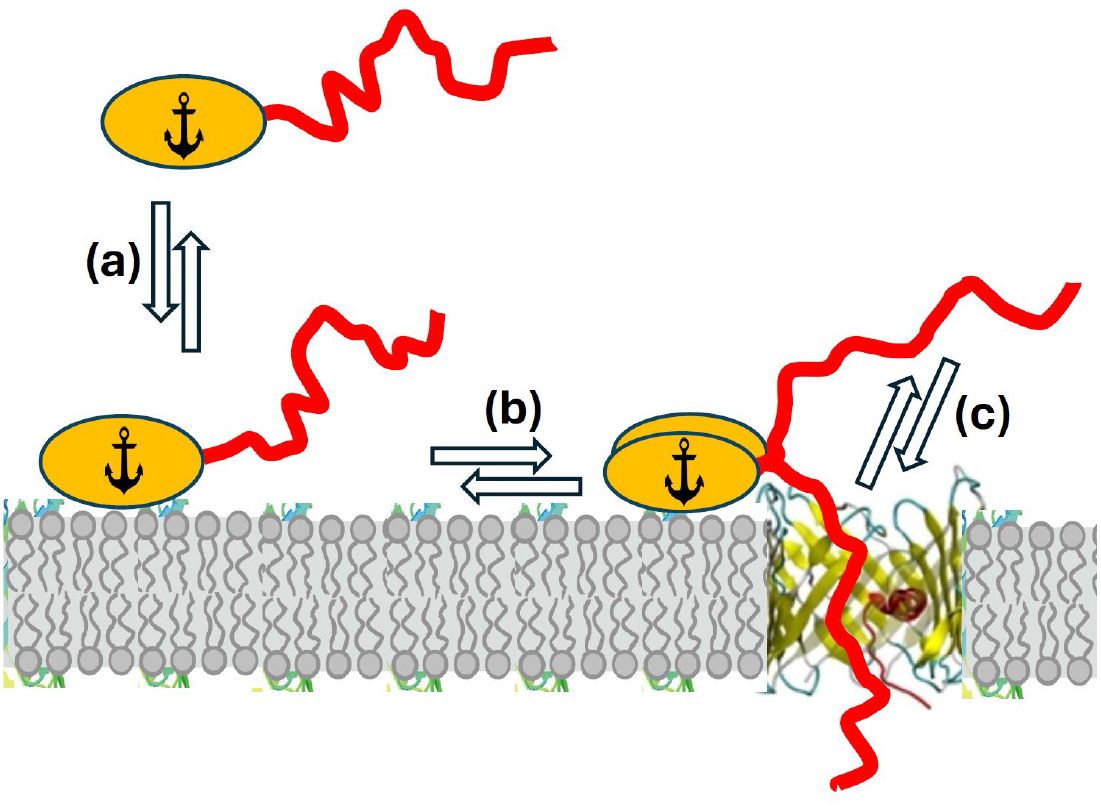
(*single column*).Model of VDAC inhibition by cytosolic peripheral proteins with charged termini. The blockage reaction consists of at least three distinct reversible steps. (a) Highly lipid-composition-dependent step of protein adsorption to the membrane surface, (b) Step of protein diffusion along the membrane surface to the VDAC vicinity, and (c) Highly voltage-dependent step of the charged terminus capture by the VDAC pore.

Encouraged by these findings, we synthesized and tested synthetic small-peptide constructs for VDAC regulation that possess the membrane-binding and channel-blocking domains. The synthesized peptides exhibited VDAC inhibition analogous to that of αSyn, particularly in terms of the characteristic time of the blockage and its exponential dependence on the applied transmembrane voltage. The on-rates of the blockage were approximately a hundred times lower, pointing to reduced accessibility of the negatively charged domains of the peptides for VDAC capture compared to αSyn, resulting in lower efficiency. However, the similarities between the synthesized peptides’ and αSyn action on VDAC allow us to conclude that our work serves as a demonstration of principle towards future development of VDAC-based mitochondrial pharmacology.

## Results and discussion

### Empirical findings on blockage kinetics and equilibria

Based on the findings briefly summarized above, we designed three peptides comprising the membrane-binding and pore-blocking domains, as shown in **Fig. 2**. For the membrane-binding domains, we have chosen the 23-residue-long hydrophobic N-terminus of hexokinase 2 (HK2p) or the 6-25 amino acids of αSyn N-terminus ^33^. For the pore-blocking domain, the acidic CTT sequence of β3-tubulin was selected. The latter choice was based on our earlier finding in experiments with recombinant tubulins ^34^ that this sequence was the most effective VDAC blocker among the tested tubulin CTTs. The rationale of selecting HK2p is that it was identified as the hexokinase 2 mitochondrial-binding domain, with a measured *K*_*d*_ of about 30 µM. ^33, 35^. Thus, using HK2p may serve not only as a membrane anchor but also as the sequence that selectively targets mitochondria. Correspondingly, the 6-25 N-terminus of αSyn has been shown to be a high-affinity membrane-binding domain with an estimated *K*_*d*_ in the µM range ^36, 37^.

**Fig. 2.**
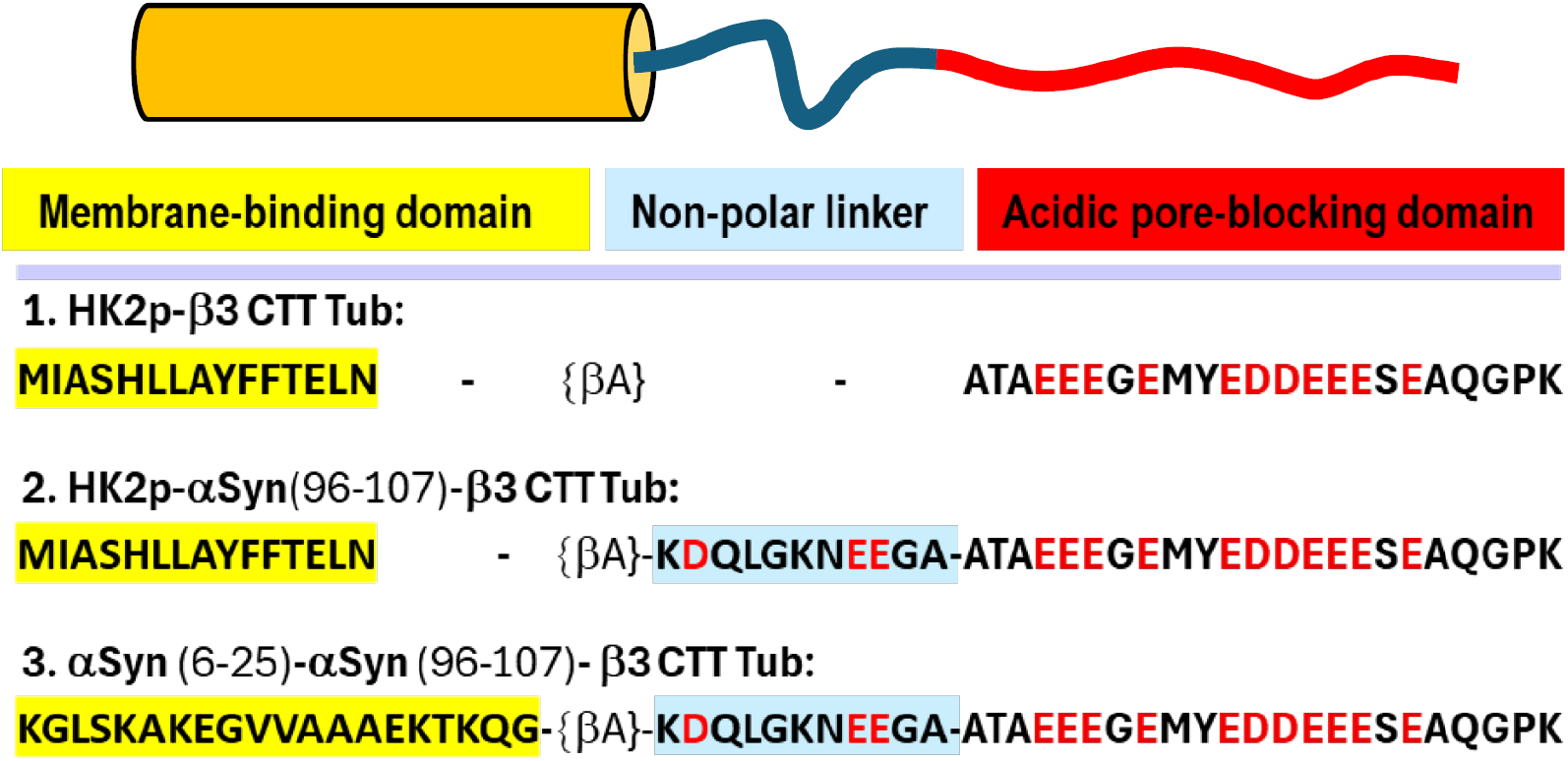
(*single column*).Structure of chimeras used in the study. The peptides that block VDAC in a controllable way were engineered using αSyn as a template of the VDAC regulator. The VDAC blocker molecule is water-soluble and possesses a three-domain structure, comprising a membrane-binding domain (yellow), affixing the blocker on the membrane surface; a linker domain (blue); and a pore-blocking acidic domain (red). The membrane-binding domains correspond to the 23-residue-long hydrophobic N-terminus of hexokinase2 (HK2p) (chimeras 1 and 2) or to the 6-25 amino acids of αSyn N-terminus (chimera 3). The acidic β3-tubulin CTT was chosen as the pore-blocking domain for all three chimeras, as it was the most effective VDAC blocker among the tested tubulin CTTs ^34^. Upon association with target membranes using its membrane-binding domain, a blocker “floats” on the membrane surface in a raft-like manner until it encounters the VDAC pore, where the acidic pore-blocking domain comes into action. A linker domain allows fine-tuning of the efficacy of the acidic domain in recognizing the VDAC pore.

**Fig. 3** provides a concise summary of the raw data on VDAC blockage by the three chimeras, compared with αSyn and β3-tubulin, along with the corresponding blockage time histograms shown on the right. It is seen that the chimeras induce characteristic transient blockades of the VDAC conductance with amplitude and average durations similar to those of αSyn, thus providing a proof-of-principle demonstration. At that, appreciable blockage requires application of the constructs in the micromolar rather than nanomolar concentration range.

**Fig. 3.**
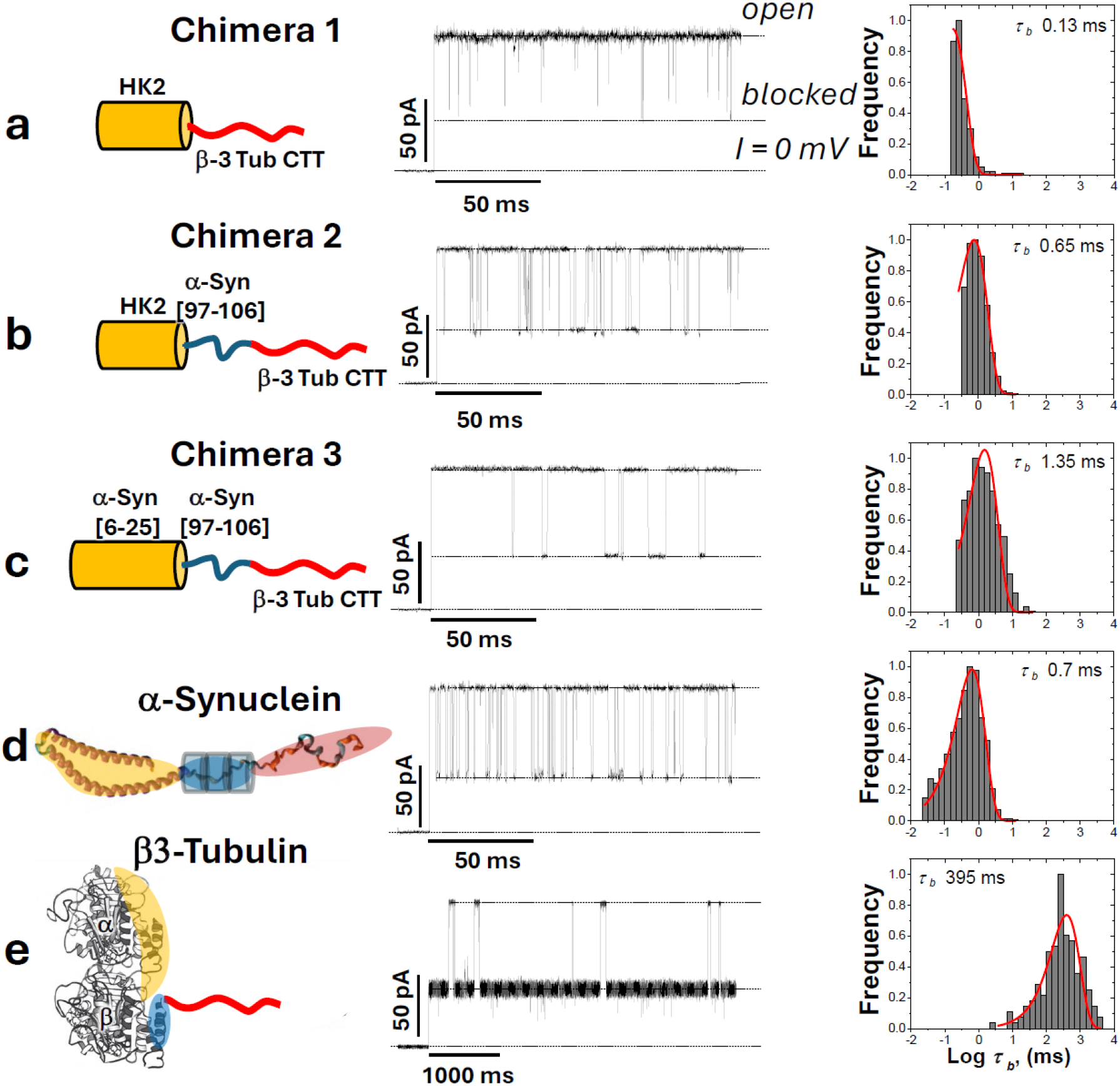
(*double column*).Sample traces of VDAC blockage by the three chimeras in comparison with two endogenous VDAC blockers, dimeric tubulin and αSyn, and the corresponding distributions of the blockage times. *Left column*: (a)-(c) Cartoons illustrating three blocking constructs, (d) αSyn, and (e) recombinant tubulin β3. Color coding indicates three specific blocker domains: the membrane-binding domain (yellow), the linker domain (blue), and the pore-blocking domain (red). *Middle column:* representative traces of current through single VDAC channels recorded in the presence of 5 µM of chimera 1 and chimera 2 constructs ((a) and (b), respectively), 10 µM of chimera 3 construct (c), 10 nM of αSyn (d), and 6 nM of recombinant β3-tubulin construct (e). All records were obtained at -30 mV of the applied voltage; the membrane-bathing solutions contained 1 M KCl buffered with 5 mM HEPES at pH 7.4. Dashed lines indicate VDAC open and blocked states, and zero current. For clarity of presentation, the current records were inverted and smoothed with a 5-point boxcar digital filter using pClamp v.10.3. *Right column*: corresponding log-binned distributions of blockage times, τ_*b*_, calculated from statistical analysis of current records, fragments of which are shown in the middle. Red solid lines are the single exponential fits with characteristic time τ_*off*_ = = <τ_*b*_ > indicated on each histogram. The total number of events represented by histograms in (a), (b), (c), (d), and (e) are 724, 2410, 740, 156, and 2398, respectively.

The results of statistical analysis of the on- and off-rates shown in **Figs. 4a** and **4c** demonstrate an exponential dependence of both rates on the applied voltage. The resulting equilibrium constants for the chimeras and the cytosolic proteins, *k*_*eq*_ *= k*_*on*_ *\**τ_*off*_ (**Fig. 4b)**, are, therefore, also exponential in voltage. The sensitivities of the equilibrium constants to voltage are summarized in **Table 1** in terms of the effective gating charge.

**Table 1.** Effective “gating charges” of VDAC blockage by three chimeric constructs, calculated from the measured voltage dependences for *k*_*on*_, *t*_*off*_, and *k*_*eq*_ shown in Fig. 4 through their fitting as described in the Fig. 4 caption, in comparison with the values calculated earlier for αSyn ^22^ and β3-CTT tubulin construct ^34^. Gating charge values are given in the units of elementary charge as determined from the mean of 3-5 independent experiments ± S.D.

| Chimera/<br>Protein | $q_{on}$ | $q_{off}$ | $q_{eq}$ |
| --- | --- | --- | --- |
| Chimera 1 | $2.77^{+0.09}_{-0.09}$ | $3.61^{+0.06}_{-0.07}$ | $6.31^{+0.14}_{-0.14}$ |
| Chimera 2 | $3.00^{+0.14}_{-0.12}$ | $3.99^{+0.12}_{-0.11}$ | $7.23^{+0.14}_{-0.15}$ |
| Chimera 3 | $2.22^{+0.29}_{-0.23}$ | $6.39^{+0.35}_{-0.44}$ | $8.42^{+0.67}_{-0.83}$ |
| αSynuclein | $3.49^{+0.12}_{-0.12}$ | $6.60^{+0.31}_{-0.35}$ | $10.02^{+0.33}_{-0.41}$ |
| β3-Tubulin | $4.12^{+0.13}_{-0.11}$ | $9.91^{+0.22}_{-0.17}$ | $13.15^{+0.26}_{-0.57}$ |

**Fig. 4.**
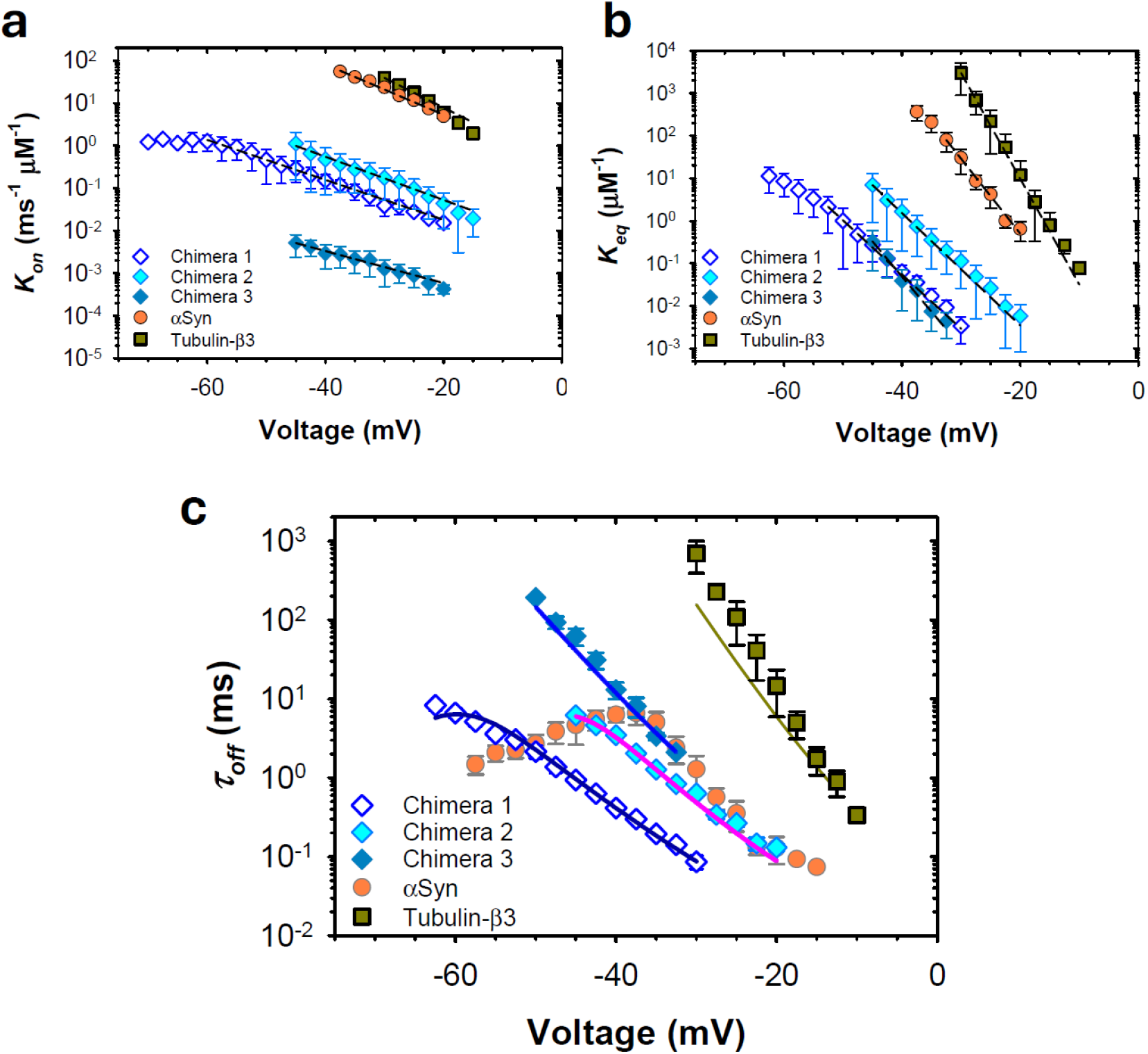
(*double column*).Kinetics of the interaction of the three constructs and two endogenous blockers with VDAC. Voltage-dependences of the on-rate, *k*_*on*_, (a), equilibrium constants, *k*_*eq*_ *= k*_*on*_ *\**τ_*off*,_ (b), and blockage time, τ_*off*_, (**c**), obtained for chimera 1, chimera 2, chimera 3 constructs in comparison with αSyn and β3-tubulin. The on-rate and equilibrium constants of VDAC blockage by three constructs are exponentially dependent on the applied voltage. The dashed straight lines in panels (a) and (b) are fits to *k*_*on*_ (*V*) = *k*_*on*_(0)exp[|*V*|*q*_*on*_/(*k*_*B*_*T*)], *k*_*eq*_ (*V*) = *k*_*eq*_(0)exp[|*V*|*q*_*eq*_/(*k*_*B*_*T*)]. The fits to *t*_*off*_ (*V*) = *t*_*off*_(0)exp[|*V*|*q*_*off*_/(*k*_*B*_*T*)] in panel (c) are not shown to not obscure the predictions of the modeling indicated by the solid lines (see the text). Here, *V* is the applied voltage, and *q* is the effective “gating charge”, summarized in Table 1, with *e, k*_*B*_, and *T* having their usual meaning of the elementary charge, Boltzmann constant, and absolute temperature, respectively. The increase of blockage time with absolute amplitude of the applied voltage (c) corresponds to the pore blockage regime; the decrease of blockage time with voltage amplitude corresponds to the regime of αSyn translocation through the pore ^38^. The solid lines in (c) are fits obtained from the energy landscape modeling. Each data point is a mean of 3-5 independent experiments ±S.D. (error bars).

### Energy landscape modeling

To understand the physical mechanism behind the observed dynamic behavior of each chimera, we turn to energy landscape modeling, a well-established technique for quantifying the forces acting on a trapped polymer chain. Energy landscape modeling involves constructing a model *U*(*x*) of the one-dimensional energy profile experienced by the chain as a function of the distance *x* it has moved beyond the constriction of the channel ^39-41^. We will use a model proposed by us earlier ^23, 42, 43^, in which one end of the profile (*x* = 0) corresponds to retraction of the polypeptide strand into the same reservoir from which it was captured; the other end (*x* = *L*, for total polypeptide length *L*), represents translocation of the entire polypeptide strand into the opposite reservoir ^38^. The average escape time of the polypeptide, calculated as the mean first passage time to either end of the profile, corresponds to *τ*_*off*_ observed in the experiments. The expression is, for *Ũ* (*x*) = *U*(*x*)/*k*_*B*_*T* and a starting position *x*_0_,

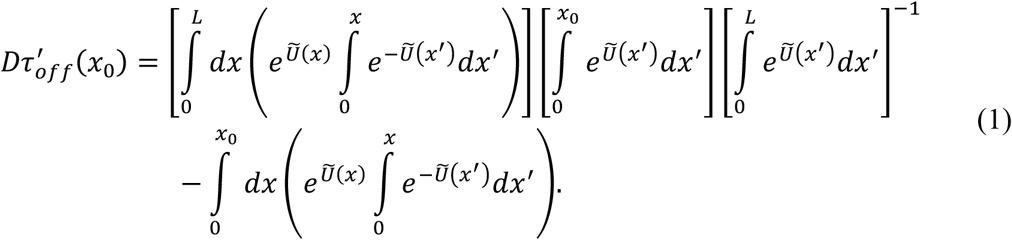

The escape times to be compared to experiments are averaged over a distribution *P*(*x*_0_) of initial positions between 5 and 20 nm, weighted by a Boltzmann distribution, *i*.*e*. 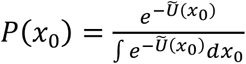, to approximate the equilibrium distribution of occupancy. Thus,

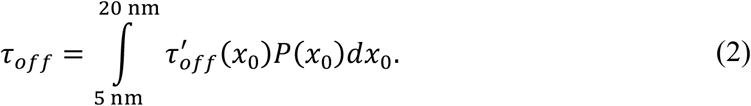

Contributing elements of the free energy profile are as follows.

i. Entropic forces from confinement of the polypeptide to the pore:

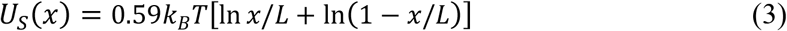
ii. Electrostatic forces arising from the interaction of the applied electric field with the linear charge density *σ*_*bare*_ from the charged residues, accounting for electroosmotic effects through the parameters *m*_*EOF*_ and *b*_*EOF*_:

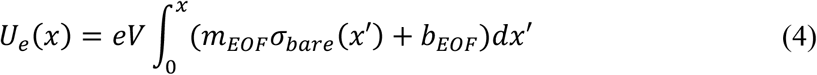
iii. Binding energy of the polypeptide to the surface, modeled as a sigmoidal barrier:

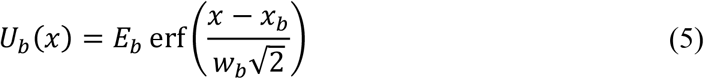

The full free energy profile is *U*(*x*) = *U*_*S*_(*x*) + *U*_*e*_(*x*) + *U*_*b*_(*x*). Of particular interest are the strength of the binding interaction (*E*_*b*_), and the position of the residue at which the binding retards the motion of the polypeptide (*x*_*b*_).

**Fig. 5** shows the results of the energy landscape model after optimizing the model parameters based on the observed *τ*_*off*_, with the solid lines in Fig. 4c indicating optimized energy landscape model predictions. *E*_*b*_ clearly corresponds to the height of the energy barrier in the absence of electrostatic forces; as electrostatic forces increase, a metastable energy well is created around *x* ≈ 8 nm for each construct. In the voltage range shown, the depth of this well increases, leading to the observed exponential increase in *τ*_*off*_. Importantly, the energy landscape models allow identification of the residues at the edge of the binding region (vertical arrows). This allows us to create a structural model of each chimera’s interaction with the VDAC channel (**Fig. 6**).

**Fig. 5.**
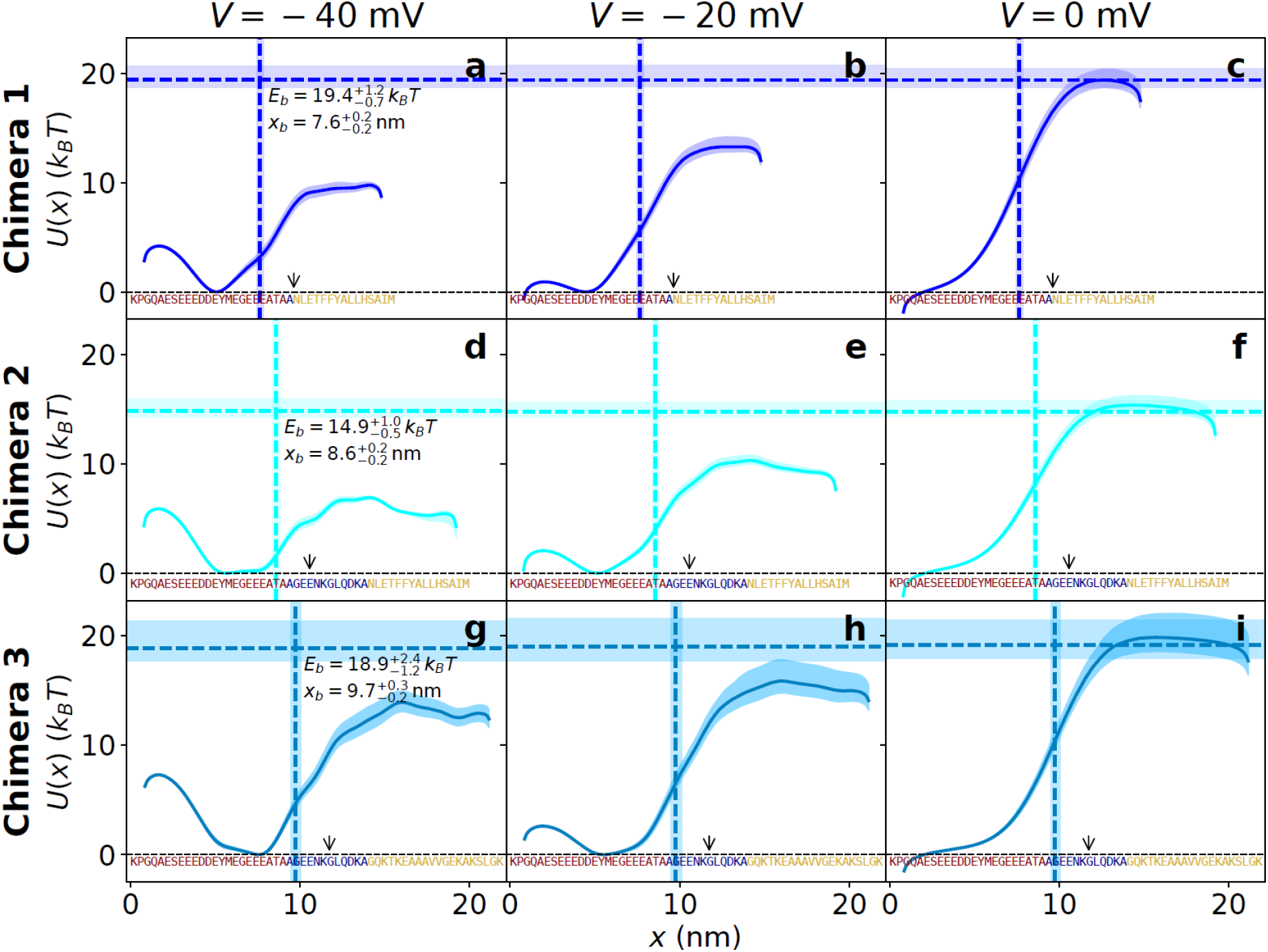
(*double column*).Energy landscape modeling. (a-c, d-f, g-i) Optimized free energy profiles for each of chimeras 1, 2, and 3, at *V* = −40, −20, and 0 mV, respectively. The horizontal axis is the length of polypeptide that has moved through the nanopore; zero thus represents the last residue of the CTT. Shading shows the 68 % confidence interval. The vertical dashed line (68% CI shaded) indicates the optimized barrier position, which is at the membrane center; the arrows show the estimated position of the membrane surface.

**Fig. 6.**
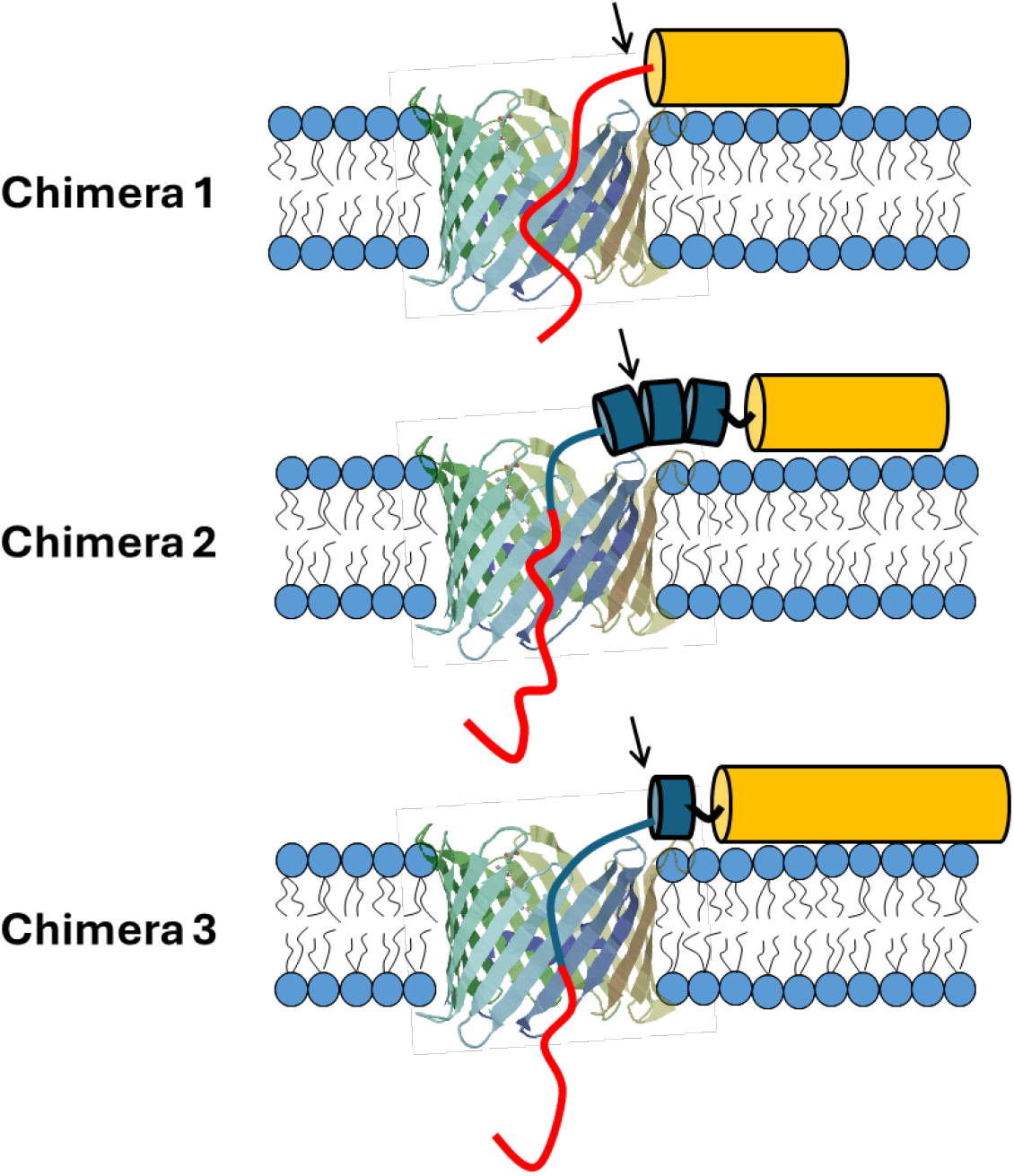
(*single column*)Visual representation of one of the energy landscape modeling results. Black arrows correspond to those in Fig. 5 and show the location of the last residue on the surface near the lip of the VDAC channel.

When the CTT of chimera 1 is captured in the VDAC channel, the CTT first residue is at the end of the membrane-anchoring domain at the membrane surface. Chimera 2, by contrast, was designed to include a non-polar, flexible linker separating the membrane from the CTT. The purpose of this linker was to allow the CTT to penetrate deeper into the VDAC channel, leading to longer-lived states ^44^. Interestingly, this linker was only partially effective. Fig. 5 shows that the penetration into the channel increases by only about 2 residues, enough to draw an additional charge into the channel and increase the effective gating charge (Table 1); however, the entire linker is not drawn into the pore. This may be due to the unintended tendency of residues in the linker region to form a helix adjacent to the membrane, or simply to the unexpected association of the linker region with the membrane (Fig. 6). The reduced energy barrier to unbinding may also indicate destabilization of the hexokinase-derived membrane-binding domain.

Chimera 3 was designed similarly to chimera 2, with the flexible linker, but the membrane-binding domain is derived from αSyn. Energy landscape modeling suggests that the binding affinity of the αSyn-derived binding domain is quite similar to the hexokinase-derived binding domains of chimeras 1 and 2 (*K*_*d*_ ≈ 30 *μM*) ^45^; this result is consistent with NMR estimates of the membrane binding affinity of the isolated αSyn-derived binding domain with various lipid compositions ^37^. Energy landscape modeling further shows a deeper average penetration of CTT into the VDAC channel, leading to an increased *τ*_*off*_. This deeper penetration appears to come at a cost, however; Fig. 4 shows a significantly reduced capture rate *k*_*on*_. Previously, it was shown for αSyn that disordered conformations, in which the CTT is not close to the VDAC channel entrance, have a characteristic combination of high *τ*_*off*_ but low *k*_*on*_^44^. To test whether such conformations may be present with Chimera 3, we analyzed the distributions of capture events under asymmetric salt conditions. As shown previously, these conditions change the apparent conductance of the channel (defined as the ratio of the current through the blocked channel to the applied voltage) depending on the average charge of the polypeptide strand occupying the lumen ^38, 46^, and can aid in distinguishing different binding conformations ^44, 47^. Distributions are shown in **Fig. 7**. Chimera 3 clearly shows multiple populations of capture events, suggesting significant conformational heterogeneity. We attribute disorder in the linker region of chimera 3 (Fig. 6, lower panel) as the likely cause of these observations. By contrast, chimeras 1 and 2 show homogeneous distributions.

**Fig. 7.**
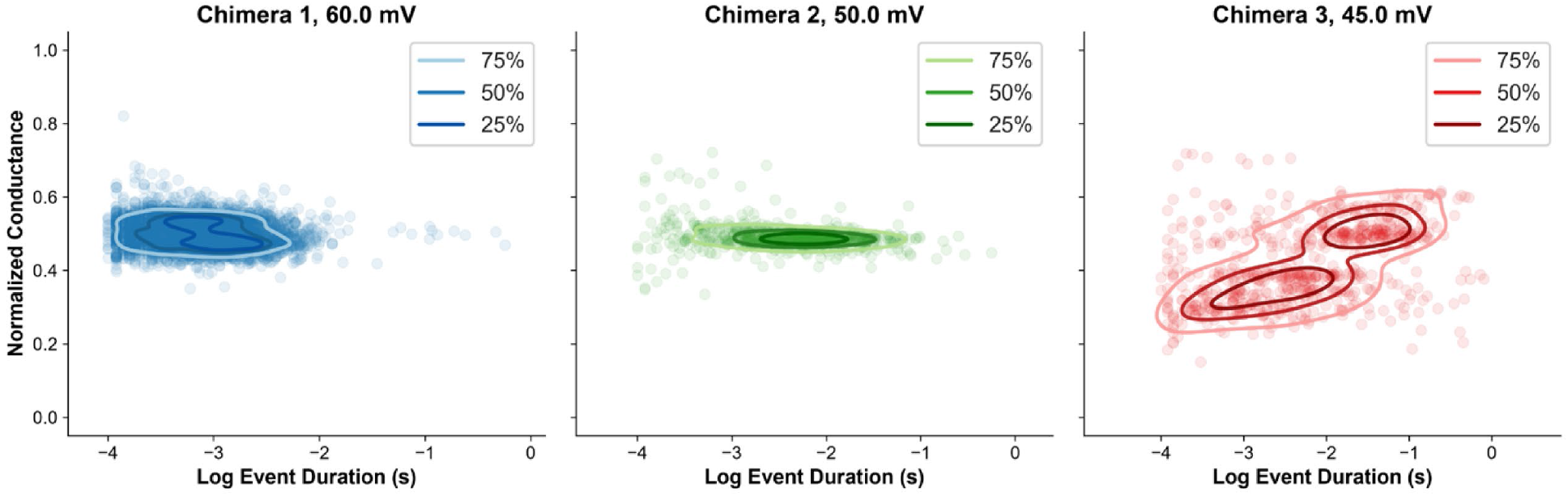
(*double column*).Distributions of apparent conductance (normalized to the open state conductance) and duration of individual capture events of the chimeras into VDAC under asymmetric (1.0 M KCl *cis*, 0.2 M KCl *trans*) salt conditions. Density contours at the 25 %, 50 %, and 75 % levels are shown to guide the eye.

The equilibrium constant (Fig. 4b) combines both *τ*_*off*_ and *k*_*on*_ and thus provides a combined measure of each construct’s effectiveness in blocking VDAC. Chimera 2 is the most efficient; the helical hexokinase-derived binding domain pins the CTT near the membrane surface, while the flexible linker enables greater penetration compared to chimera 1. Chimera 3 shows higher penetration still but its significantly lower capture rate leads to lower overall effectiveness. Remarkably, αSyn has a much higher equilibrium constant than any of the constructs, presumably due to its extensive membrane-binding domain that displays significant membrane binding even below 0.1 µM ^22, 48^, and a favorable binding conformation that constrains the CTT near the VDAC entrance ^44^. Tubulin is yet more effective; as previously shown, tubulin binds to the membrane with a conformation that anchors the CTT in a way that it is easily captured by the VDAC pore ^34, 49^, leading to high capture rates and highly efficient blockage. Together, these results suggest that a surface density-normalized (rather than solution concentration-normalized) equilibrium constant may better characterize blockage efficiency, with the important caveat that surface density is experimentally much less accessible.

The goal of this work was to design a peptide construct with high VDAC-blocking efficiency to establish a prototype VDAC inhibitor for the future development of VDAC pharmacological tools. VDAC has recently emerged as an important anticancer target. Our results demonstrate that it is possible to achieve a fine regulation of VDAC permeability by rationally designed peptides, thus opening a new avenue in the search for VDAC therapeutics. The advantage of using such a peptide construct is that none of the three chimeras studied here can translocate through VDAC, as evidenced by the continuous exponential increase in *t*_*off*_ even at an applied voltage as high as 60 mV (Fig. 4c). This suggests restricting their action exclusively to the outer membrane and not targeting the inner membrane with electron transport chain complexes, in contrast with αSyn behavior, which translocates through VDAC, as evidenced by a decrease in *t*_*off*_ at |*V*| > ∼40 mV (Fig. 4c), and is known to target both mitochondrial membranes in cells ^50^. The next immediate step is to find a more potent and less toxic peptide for the appropriate membrane-binding domain instead of using the HK2p (see related discussion in ^33^). We expect the synthetic peptide to reduce ATP/ADP permeability through VDAC and increase Ca^2+^ uptake, based on available data on αSyn and tubulin interactions with VDAC ^24^. Increased mitochondrial Ca^2+^ uptake can trigger apoptosis in cancer cells. However, it is still too early to predict the exact outcome of using such peptides in live cells because many known players – cytosolic and mitochondrial proteins – are involved in complexation with VDAC, thereby affecting multiple crucial cellular pathways (see, e.g., the recent reviews ^51, 52^. Future studies are needed to confirm the cellular mechanism and the potential of the proposed designed peptides as anticancer drugs.

## Conclusions

We have demonstrated that artificial, chimeric peptides derived from known natural VDAC blockers ^21, 22^ also exhibit VDAC-blocking activity. These chimeras behave in remarkably similar ways despite variations in linker length and membrane-binding domains. The differences among the three chimeras, native αSyn, and recombinant β3-tubulin highlight the design principles that lead to highly efficient VDAC blockage, specifically (i) maximizing the channel penetration of the negatively charged disordered tail, (ii) ensuring that the tail is constrained by the membrane binding domain to be near the channel entrance, and (iii) minimizing unwanted secondary structure formation in the linking region and its interactions with the membrane.

## Experimental section

### Protein purification

VDAC was isolated from frozen mitochondrial fractions of rat liver that were a generous gift of Dr. Marco Colombini (University of Maryland, College Park, USA) following the standard method ^53^ and purified on hydroxyapatite/celite (2:1) column as previously described ^54^. VDAC purified from the mitochondrial fraction of the liver contains all three isoforms with VDAC1 being the predominant one (∼ 80% of total VDACs) ^55^. An engineered recombinant tubulin dimer constructed of yeast α/ß tubulin heterodimer with human carboxy-terminal tail b3(TUBB3), was generated, isolated, and purified as previously described ^34, 56^. Recombinant αSyn was a kind gift from Dr. Jennifer Lee (NHLBI, NIH). αSyn was expressed, purified, and characterized as described previously ^57^.

### Synthetic peptides

All three peptides were synthesized by the Research Central Facility for Biotechnology Resources, FDA.

### Channel Reconstitution

Planar bilayer membranes were formed from lipid monolayers across a ∼ 70-90 μm orifice in a 15 μm Teflon film separating two ∼ 1.2 mL reconstitution cell compartments, as described previously ^58^. A 5 mg/mL solution of diphytanoyl-phosphatidylcholine (DPhPC) (Avanti Polar Lipids, Alabaster, AL) in pentane was used for bilayer formation. Channel insertion was achieved by adding purified VDAC in a 2.5% Triton X-100 solution to the aqueous phase of 1 M KCl buffered with 5 mM HEPES at pH 7.4 in the *cis* compartment while stirring. Potential is defined as positive when it is greater at the side of VDAC addition (*cis*). Tubulin was added to the membrane-bathing solutions in the *cis* compartment after VDAC reconstitution and collection of control recordings of the channel current in tubulin-free solution. Conductance measurements were performed using an Axopatch 200B amplifier (Axon Instruments, Inc., Foster City, CA) in the voltage clamp mode. Data were filtered using a low-pass 8-pole Butterworth filter (Model 900, Frequency Devices, Inc., Haverhill, MA) at 15 kHz and a low-pass Bessel filter at 10 kHz and directly saved into computer memory with a sampling frequency of 50 kHz. The effective time resolution of the setup was approximately 200 μs. For data analysis by Clampfit 10.3, no digital filtering was applied. Individual events of current blockages were discriminated, and kinetic parameters were obtained by fitting logarithmic single-, double-, or triple-exponentials to logarithmically binned histograms ^59^. All lifetime histograms used 10 bins per decade. The number of blocking events for each analyzed current fragment was in the range from 250 to 2500. Four different logarithmic probability fits were generated using different fitting algorithms, and the mean and standard deviation of the fitted time constants were used as the mean and standard deviation for the characteristic open and blockage times.

### Energy landscape modeling

As described previously ^23, 42, 43^, the energy landscape model is constructed as the sum of three terms: 1. An energy barrier to unbinding, modeled as a sigmoid parameterized by height, position, and width. The height represents the strength of binding, while the position identifies the residue in the center of the VDAC channel when the CTT’s motion is arrested by the binding domain. A surface position, representing the residue at the surface of the membrane facing the *cis* reservoir, is derived from the barrier position as 5 residues closer to the N-terminus, based on a length per residue of about 0.4 nm ^60^ and a ≈ 2 nm distance from the membrane surface to the center of the channel ^61^. 2. Electrostatic energy arises from interaction of charged residues with the electric field but is counteracted by electroosmotic flow; this term is parameterized by the slope *m*_*EOF*_ and intercept *b*_*EOF*_, which define the relationship between electroosmotic flow and polypeptide charge. The effective charge density is smoothed by the characteristic width of the helical constriction at the center of VDAC (Hoogerheide 2020). 3. Entropic energy of confining the polypeptide in the pore, assuming the peptide is a disordered polymer. The diffusion constant is fixed at 0.52 μm^2^ s^−1^ based on previous determinations ^23^. The injection point for the first-passage problem is set to a Boltzmann distribution in the energy landscape near its minimum.

### Data fitting

Energy landscape and gating charge models were optimized using a Markov Chain Monte Carlo (MCMC) algorithm implemented in the Python software package Bumps. Uncertainties are estimated from the 68 % confidence intervals of at least 5000 samples from the posterior distributions of the parameters.

## Data availability

The data supporting the findings of this study are available from the corresponding author upon reasonable request.

## Author Contributions

All authors contributed equally to this work.

## Acknowledgements

The study was funded by the NIH grant ZIA HD000072. The contributions of the NIH authors are considered Works of the United States Government. The findings and conclusions presented in this paper are those of the authors and do not necessarily reflect the views of the National Institutes of Health or the U.S. Department of Health and Human Services. Certain commercial materials, equipment, and instruments are identified in this work to describe the experimental procedure as completely as possible. In no case does such an identification imply a recommendation or endorsement by NIST, nor does it imply that the materials, equipment, or instruments identified are necessarily the best available for the purpose. Unless otherwise noted, NIST work was funded solely by the United States Government.

## References

1. J. P. Overington, B. Al-Lazikani and A. L. Hopkins, Nat Rev Drug Discov, 2006, 5, 993–996.

2. J. J. Clare, Discov Med, 2010, 9, 253–260.

3. A. Wickenden, B. Priest and G. Erdemli, Future Medicinal Chemistry, 2012, 4, 661–679.

4. H. Wulff, P. Christophersen, P. Colussi, K. G. Chandy and V. Yarov-Yarovoy, Nat Rev Drug Discov, 2019, 18, 339–357.

5. D. C. Camerino, D. Tricarico and J. F. Desaphy, Neurotherapeutics, 2007, 4, 184–198.

6. H. Wulff and B. S. Zhorov, Chemical Reviews, 2008, 108, 1744–1773.

7. J. J. Clare, Expert Opin Inv Drug, 2010, 19, 45–62.

8. E. M. Nestorovich and S. M. Bezrukov, Chemical Reviews, 2012, 112, 6388–6430.

9. S. Bagal, A. D. Brown, P. J. Cox, K. Omoto, R. M. Owen, D. C. Pryde, B. Sidders, S. E. Skerratt, E. B. Stevens, R. I. Storer and N. A. Swain, J Med Chem, 2013, 56, 593–624.

10. A. M. Waszkielewicz, A. Gunia, N. Szkaradek, K. Sloczynska, S. Krupinska and H. Marona, Curr Med Chem, 2013, 20, 1241–1285.

11. A. L. Capatina, D. Lagos and W. J. Brackenbury, Rev Physiol Bioch P, 2022, 183, 1–43.

12. K. Kofman and M. Levin, Prog Biophys Mol Bio, 2024, 191, 25–39.

13. J. T. Varughese, S. K. Buchanan and A. S. Pitt, Cells, 2021, 10.

14. M. Colombini, Molecular and Cellular Biochemistry, 2004, 256, 107–115.

15. J. J. Lemasters and E. Holmuhamedov, Biochim Biophys Acta, 2006, 1762, 181–190.

16. A. K. S. Camara, Y. Zhou, P. C. Wen, E. Tajkhorshid and W. M. Kwok, Front Physiol, 2017, 8, 460.

17. A. Magrì, S. Reina and V. De Pinto, Front Chem, 2018, 6.

18. D. Ben-Hail, R. Begas-Shvartz, M. Shalev, A. Shteinfer-Kuzmine, A. Gruzman, S. Reina, V. De Pinto and V. Shoshan-Barmatz, J Biol Chem, 2016, 291, 24986–25003.

19. S. Reina and V. De Pinto, Curr Med Chem, 2017, 24, 4447–4469.

20. T. K. Rostovtseva, M. Queralt-Martín, W. M. Rosencrans and S. M. Bezrukov, Front Physiol, 2020,11.

21. T. K. Rostovtseva, K. L. Sheldon, E. Hassanzadeh, C. Monge, V. Saks, S. M. Bezrukov and D. L. Sackett, Proceedings of the National Academy of Sciences of the United States of America, 2008, 105, 18746–18751.

22. T. K. Rostovtseva, P. A. Gurnev, O. Protchenko, D. P. Hoogerheide, T. L. Yap, C. C. Philpott, J. C. Lee and S. M. Bezrukov, J Biol Chem, 2015, 290, 18467–18477.

23. D. P. Hoogerheide, P. A. Gurnev, T. K. Rostovtseva and S. M. Bezrukov, Proteomics, 2021, 2100060.

24. T. K. Rostovtseva, S. M. Bezrukov and D. P. Hoogerheide, Int J Mol Sci, 2021, 22.

25. G. Anderluh and J. H. Lakey, Trends in biochemical sciences, 2008, 33, 482–490.

26. T. Saetersdal, G. Greve and H. Dalen, Histochemistry, 1990, 95, 1–10.

27. M. Carre, N. Andre, G. Carles, H. Borghi, L. Brichese, C. Briand and D. Braguer, J Biol Chem, 2002, 277, 33664–33669.

28. W. W. Li, R. Yang, J. C. Guo, H. M. Ren, X. L. Zha, J. S. Cheng and D. F. Cai, Neuroreport, 2007, 18, 1543–1546.

29. M. Robotta, H. R. Gerding, A. Vogel, K. Hauser, S. Schildknecht, C. Karreman, M. Leist, V. Subramaniam and M. Drescher, Chembiochem, 2014, 15, 2499–2502.

30. T. K. Rostovtseva, P. A. Gurnev, M. Y. Chen and S. M. Bezrukov, J. Biol. Chem., 2012, 287, 29589–29598.

31. T. K. Rostovtseva, D. P. Hoogerheide, A. Rovini and S. M. Bezrukov, in Biol Med Phys Biomed, 2017, 10.1007/978-3-319-55539-3_8, ch. Chapter 8, pp. 185–215.

32. D. Jacobs, D. P. Hoogerheide, A. Rovini, Z. Jiang, J. C. Lee, T. K. Rostovtseva and S. M. Bezrukov, Sci Rep, 2019, 9, 4580.

33. M. Rajendran, M. Queralt-Martin, P. A. Gurnev, W. M. Rosencrans, A. Rovini, D. Jacobs, K. Abrantes, D. P. Hoogerheide, S. M. Bezrukov and T. K. Rostovtseva, Cell Mol Life Sci, 2022, 79, 368.

34. T. K. Rostovtseva, P. A. Gurnev, D. P. Hoogerheide, A. Rovini, M. Sirajuddin and S. M. Bezrukov, J Biol Chem, 2018, 293, 10949–10962.

35. N. Bryan and K. P. Raisch, Biosci Rep, 2015, 35.

36. G. Fusco, A. De Simone, T. Gopinath, V. Vostrikov, M. Vendruscolo, C. M. Dobson and G. Veglia, Nat Commun, 2014, 5, 3827.

37. W. K. Man, B. Tahirbegi, M. D. Vrettas, S. Preet, L. M. Ying, M. Vendruscolo, A. De Simone and G. Fusco, Nat Commun, 2021, 12.

38. D. P. Hoogerheide, P. A. Gurnev, T. K. Rostovtseva and S. M. Bezrukov, Biophys J, 2018, 114, 772–776.

39. W. Sung and P. J. Park, Phys Rev Lett, 1996, 77, 783–786.

40. M. Muthukumar, J Chem Phys, 1999, 111, 10371–10374.

41. M. Muthukumar, Polymer translocation, CRC Press, 2011.

42. D. P. Hoogerheide, P. A. Gurnev, T. K. Rostovtseva and S. M. Bezrukov, Nanoscale, 2017, 9, 183–192.

43. D. P. Hoogerheide, J Res Natl Inst Stan, 2020, 125.

44. D. P. Hoogerheide, T. K. Rostovtseva and S. M. Bezrukov, Biochimica Et Biophysica Acta-Biomembranes, 2021, 1863.

45. M. Rajendran, M. Queralt-Martín, P. A. Gurnev, W. M. Rosencrans, A. Rovini, D. Jacobs, K. Abrantes, D. P. Hoogerheide, S. M. Bezrukov and T. K. Rostovtseva, Cellular and Molecular Life Sciences, 2022, 79.

46. D. Hoogerheide, P. Gurnev, T. Rostovtseva and S. Bezrukov, Nanoscale, 2020, 12, 11070.

47. D. P. Hoogerheide, T. K. Rostovtseva, D. Jacobs, P. A. Gurnev and S. M. Bezrukov, ACS Nano, 2021, 15, 989–1001.

48. D. P. Hoogerheide and S. M. Bezrukov, J Phys Chem Lett, 2026, 17, 3917–3923.

49. D. P. Hoogerheide, S. Y. Noskov, D. Jacobs, L. Bergdoll, V. Silin, D. L. Worcester, J. Abramson, H. Nanda, T. K. Rostovtseva and S. M. Bezrukov, Proceedings of the National Academy of Sciences of the United States of America, 2017, 114, E3622–E3631.

50. A. Rovini, P. A. Gurnev, A. Beilina, M. Queralt-Martin, W. Rosencrans, M. R. Cookson, S. M. Bezrukov and T. K. Rostovtseva, Cell Mol Life Sci, 2020, 77, 3611–3626.

51. N. V. Belosludtseva, M. V. Dubinin and K. N. Belosludtsev, Biochemistry (Mosc), 2024, 89, 1061–1078.

52. M. Rajendran, S. M. Bezrukov and T. K. Rostovtseva, Int J Mol Sci, 2026, 27.

53. E. Blachly-Dyson, S. Peng, M. Colombini and M. Forte, Science, 1990, 247, 1233–1236.

54. F. Palmieri and V. De Pinto, J Bioenerg Biomembr, 1989, 21, 417–425.

55. T. Yamamoto, A. Yamada, M. Watanabe, Y. Yoshimura, N. Yamazaki, Y. Yoshimura, T. Yamauchi, M. Kataoka, T. Nagata, H. Terada and Y. Shinohara, J Proteome Res, 2006, 5, 3336–3344.

56. M. Sirajuddin, L. M. Rice and R. D. Vale, Nat Cell Biol, 2014, 16, 335–344.

57. C. M. Pfefferkorn and J. C. Lee, J Phys Chem B, 2010, 114, 4615–4622.

58. T. K. Rostovtseva, N. Kazemi, M. Weinrich and S. M. Bezrukov, J Biol Chem, 2006, 281, 37496–37506.

59. F. J. Sigworth and S. M. Sine, Biophys J, 1987, 52, 1047–1054.

60. R. K. Ainavarapu, J. Brujic, H. H. Huang, A. P. Wiita, H. Lu, L. W. Li, K. A. Walther, M. Carrion-Vazquez, H. B. Li and J. M. Fernandez, Biophys J, 2007, 92, 225–233.

61. R. Ujwal, D. Cascio, J. P. Colletier, S. Faham, J. Zhang, L. Toro, P. P. Ping and J. Abramson, Proceedings of the National Academy of Sciences of the United States of America, 2008, 105, 17742–17747.

